# Functional interrogation of neuronal connections by chemoptogenetic presynaptic ablation

**DOI:** 10.1101/2025.04.04.647277

**Authors:** Hariom Sharma, Madalina A. Robea, Noel H. McGrory, Daniel C. Bazan, Edward A. Burton, Harold A. Burgess

**Affiliations:** Division of Developmental Biology, Eunice Kennedy Shriver National Institute of Child Health and Human Development, Bethesda, MD, USA; Pittsburgh Institute of Neurodegenerative Diseases, University of Pittsburgh, Pittsburgh, PA, USA; Department of Neurology, University of Pittsburgh, Pittsburgh, PA, USA; Geriatric Research, Education, and Clinical Center, Pittsburgh VA Healthcare System, Pittsburgh, PA, USA

**Keywords:** ablation, synapse, photosensitizer, dL5**, escape behavior, raphe, Mauthner cell, prepulse inhibition, zebrafish

## Abstract

Most neurons are embedded in multiple circuits, with signaling to distinct postsynaptic partners playing functionally different roles. The function of specific connections can be interrogated using synaptically localized optogenetic effectors, however these tools are often experimentally difficult to validate or produce paradoxical outcomes. We have developed a system for photoablation of synaptic connections originating from genetically defined neurons, based on presynaptic localization of the fluorogen activating protein dL5** that acts as a photosensitizer when bound to a cell-permeable dye. Using the well mapped zebrafish escape circuit as a readout, we first show that cytoplasmically expressed dL5** enables efficient spatially targeted neuronal ablation using near infra-red light. We then demonstrate that spatially patterned illumination of presynaptically localized dL5** can effectively disconnect neurons from selected downstream partners, producing precise behavioral deficits. This technique should be applicable to almost any genetically tractable neuronal circuit, enabling precise manipulation of functional connectivity within the nervous system.

## Introduction

Genetically encoded proteins that provide experimental control over neuronal activity have revolutionized neuroscience, revealing how specific neuronal cell types contribute to circuit function and behavior. These tools utilize gene regulatory elements to drive selective expression in cell types of interest, but the complexity of neuronal ramifications can make it difficult to resolve the function of downstream connections. For example, in the Drosophila brain, neurons have a median of 13 post- synaptic partners but the CT1 visual interneuron connects to 6329 downstream targets^1^. Thus, even manipulations that achieve exquisite cell-type specificity are likely to influence multiple downstream postsynaptic partners.

To restrict experimental manipulations to desired output pathways, effector proteins can be fused to synaptic localization signals, allowing specificity through spatially controlled delivery of an activating chemical or light^2–4^. For example stimulation of channelrhodopsin in axons can restrict activation to a specific output pathway^5,6^, while presynaptic expression of optogenetic inhibitors can selectively suppress transmission to downstream neurons^7,8^. These methods are ideally suited to the optically transparent larval zebrafish system but would require animals to be immobilized under a microscope to allow for spatially patterned illumination precluding their use in freely swimming assays.

Fusion of a genetically encoded photosensitizer, miniSOG to vesicle release proteins was previously used to achieve long-lasting synaptic inactivation in *C. elegans*^9^. MiniSOG generates singlet oxygen by binding flavin mononucleotide, which is ubiquitously present in cells. Exposure to 480 nm light led to a rapid reduction in synaptic currents that persisted for several hours after discontinuation of illumination. In this system, suppression is likely due to local destruction of vesicle proteins, with recovery occurring as new components are trafficked to the synapse. We were inspired by this system to develop an analogous method for synaptic ablation in zebrafish that would enable behavioral experiments in larvae after removal of selected synaptic connections.

Fluorogen activating protein dL5** (herein dL5) is a synthetic fusion of two single-chain antibody subunits that binds with high affinity to an iodine-substituted malachite green fluorogen (MG-2I). The complex generates singlet oxygen at high quantum yield when illuminated^10–12^. The dL5 system is significantly more efficient than other phototoxic alternatives^13^ and has already been successfully used to target mitochondria in zebrafish neurons, resulting in their bioenergetic collapse, depolarization and cell death^14^. Importantly, neither dL5 protein nor free MG-2I dye are photosensitizing alone. In principle one could therefore treat animals with MG-2I, ablate targets, then wash out the dye so that animals could be tested in paradigms requiring visual stimulation without danger of further damage. The complex is activated by near infra-red light (NIR, 666 nm) which has excellent tissue penetration. Moreover, dL5 efficiently lesions cells when targeted to a variety of subcellular locations, including nucleus, cytoplasm, mitochondria or cell membrane^12^. This raises the possibility of removing specific synaptic outputs from a neuron by fusing dL5 to a vesicle protein, and selectively illuminating target synapses.

In this study we confirmed that cytoplasmic dL5 can be used to lesion neurons using spatially patterned illumination. Then, by fusing dL5 with synaptophysin, we directed the trafficking of dL5 to synapses and enabled the selective ablation of output synapses from neurons of interest. We validated this method through behavioral assessment of auditory escape and prepulse inhibition, taking advantage of the detailed mechanistic understanding of these circuits. While this tool is ideally suited to the optically accessible zebrafish system, it may also be valuable in any model where it is feasible to genetically target dL5 to synapses and to deliver spatially localized illumination.

## Results

### Widefield light exposure of dL5 expressing neurons leads to efficient ablation

Previous studies in zebrafish have shown that cytoplasmic dL5 efficiently ablates zebrafish cardiac cells and that mitochondrially targeted dL5 enables neuronal ablation^12,14^. To enable flexible cell-type specific expression, we generated a transgenic line with cytoplasmic dL5 expression under control of a UAS promoter (Fig. 1A). We tested the ablation of cells expressing dL5 using Gal4 line *y252-Gal4*, because ablation of neurons labeled in this line has multiple well-characterized effects on behavior^15–17^. In *y252-Gal4*, *UAS:dL5-mCer* larvae that were treated with MG-2I for 24 h, then exposed to 550 mW/cm^2^ widefield (*i.e.* whole-body) 656 nm light for 20 min, we saw a complete loss of mCer expressing cells at 24 h (Fig. 1B, Supp. Fig. 1A). Both MG-2I treatment and intense illumination were required for ablation (Fig. 1B). At 4 hours after light exposure, mCer-expressing cells remained intact – demonstrating that loss of fluorescence is not due to photobleaching – but by 16 hours, near-total loss was observed (Fig. 1C). For wild-type embryos not expressing dL5, MG-2I treatment was non-toxic, including in embryos exposed to intense light, as evidenced by the presence of an inflated swimbladder at 6 days post-fertilization (dpf), which is a sensitive marker of health in zebrafish larvae (Supp. Fig. 1B). We tested different durations of light exposure and found a dose-response effect, with 20 min required for complete loss of *y252-Gal4* neurons (Supp. Fig. 1C).

**Figure 1.**
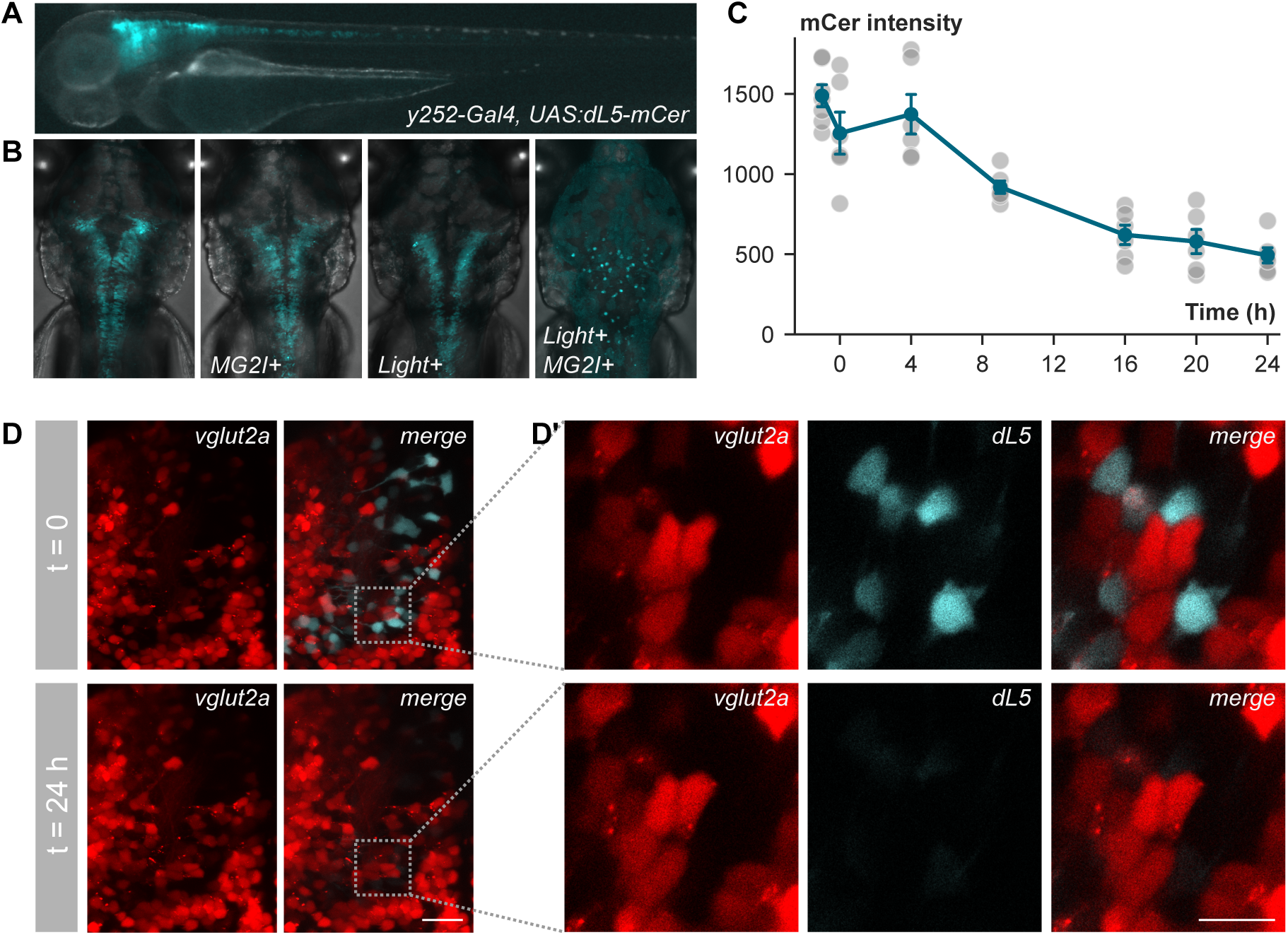
**Efficient neuronal ablation using cytoplasmic dL5** A. Transgenic *y252-Gal4, UAS:dL5-mCer* larva at 3 dpf. Merged fluorescent and visible light exposures. B. Maximum projection of confocal images of *y252-Gal4, UAS:dL5-mCer* 6 dpf larvae. Untreated (left), MG-2I only, light only and combined light and MG-2I as indicated. C. Cerulean fluorescence intensity in *y252-Gal4, UAS:dL5-mCer* immediately before and after photoablation, and at the indicated timepoints. Values are normalized to non-fluorescent sibling larvae. D. Maximum projection (D) and matched single horizontal confocal slices (D’) through part of the diencephalon of *y252:Gal4, UAS:dL5-mCer, vglut2a:DsRed* larvae immediately before exposure to NIR light (top panels) and 24 h later (bottom panels). Region in D’ is rotated from boxed area in D. Scale bars 20 µm.

Singlet oxygen has limited diffusion in a cellular environment which should help to limit collateral damage during photoablation. Indeed, in zebrafish, photostimulation of MG-2I treated mitochondrially- targeted dL5 damaged mitochondria without acutely affecting adjacent endoplasmic reticulum or Golgi apparatus^14^. However prolonged illumination of membrane-localized dL5-expressing cells can damage neighboring cells^12^. We therefore assessed the specificity of dL5 neuronal ablation by imaging cells adjacent to dL5-expressing cells in the same larva before exposure to MG-2I and NIR, and 24 h later. We focused on a diencephalic region in *y252-Gal4, UAS:dL5-mCer, vglut2a:DsRed* larvae, where *y252* expressing neurons and *vglut2a* expressing neurons are each relatively sparse and rarely overlap. There was no loss of *vglut2a:DsRed* signal in this region after photoablation (Fig. 1D), and inspection of single confocal planes demonstrates that *vglut2a:DsRed* neurons that were closely apposed to *y252- Gal4, UAS:dL5-mCer* neurons remained intact, despite the complete loss of dL5 neurons (Fig. 1D’).

Next, we checked whether photoablation with dL5 produces anticipated behavioral deficits (Fig. 2A). To test this, we utilized a previously established method of ablating neurons in the *y252-Gal4* line by driving nitroreductase (NTR) expression from a *UAS:epNTR-RFP* transgenic line^18^. NTR is a bacterial protein that converts the pro-drug metronidazole into a cell-impermeant toxin, enabling cell-specific ablation^19^. We ablated NTR expressing *y252-Gal4* neurons with metronidazole treatment from 3-4 dpf, then tested larvae at 6 dpf. As expected, we observed an increase in short-latency C-start (SLC) escapes, decrease in long-latency C-start (LLC) escapes, and reduction in prepulse inhibition (PPI) (Fig. 2B). PPI is a form of sensorimotor gating in which a weak pre-stimulus suppresses the SLC response to a subsequent intense stimulus. To assess whether dL5-based ablation reproduced these deficits, we treated larvae from 2-3 dpf with MG-2I, photoablated neurons with 20 min widefield NIR exposure, then tested larvae at 6 dpf. dL5-ablated larvae displayed the same deficits as NTR-ablated larvae: increased SLC responses, decreased LLC responses and decreased PPI (Fig. 2C). Behavioral effects after dL5 ablation were slightly weaker despite the relatively complete loss of mCer fluorescence. This may reflect differences in the perdurance of expression of dL5-mCer compared to epNTR-RFP, longer recovery time after photoablation providing more opportunity for cell replacement during development, or in the degree to which expression was variegated in each line. Nevertheless, we confirm that cytoplasmic dL5 expression coupled with MG-2I treatment and light exposure leads to a lasting loss of zebrafish neurons, that produce expected effects on behavior.

**Figure 2.**
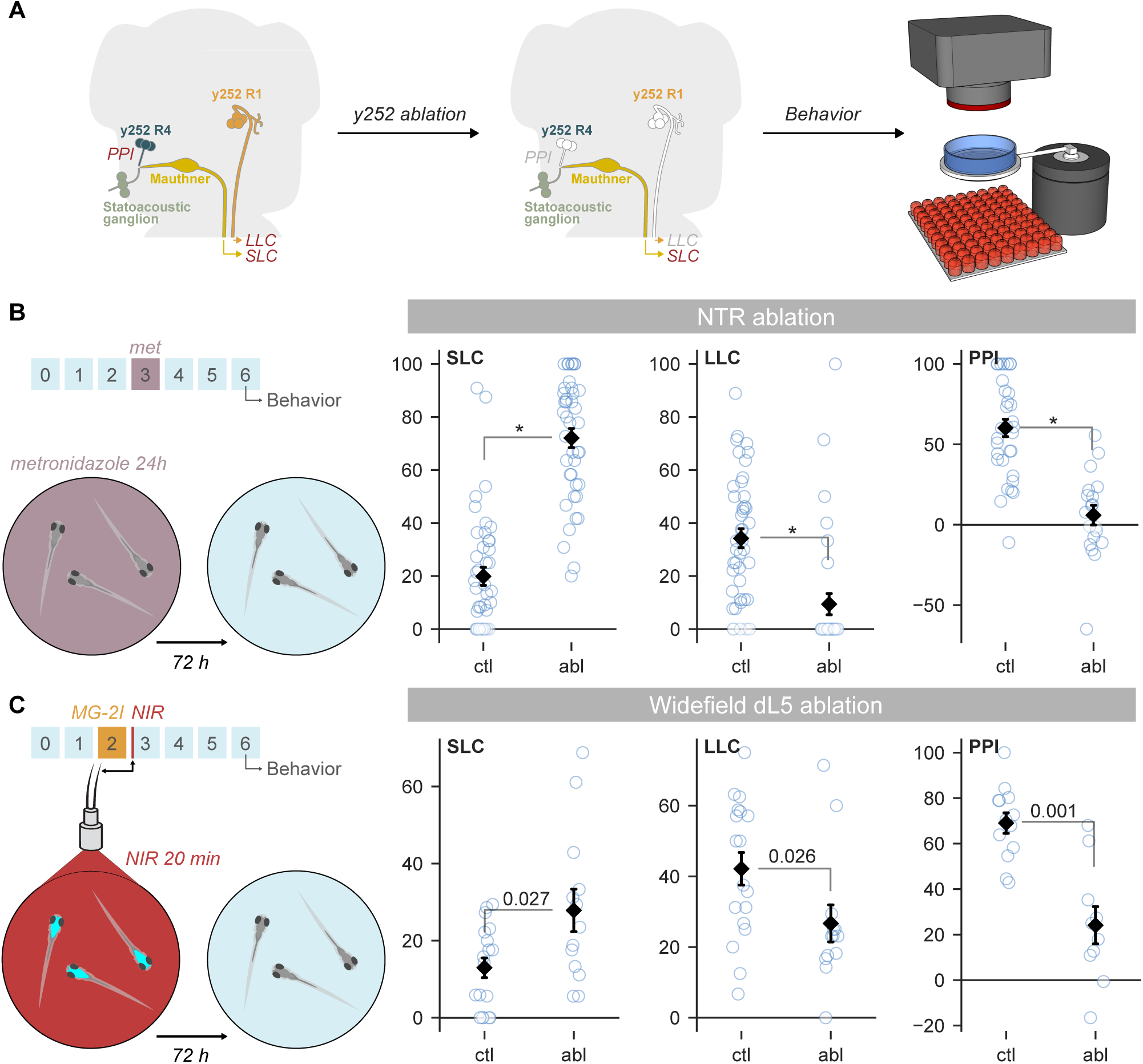
**dL5 photoablation reproduces behavioral deficits caused by NTR ablation** A. Relevant neurons for short-latency and long-latency C-start escape behavior (SLC, LLC) and prepulse inhibition (PPI). *y252-Gal4* labels rhombomere 1 (R1) neurons required for LLC responses and rhombomere 4 (R4) neurons that mediate PPI. After ablation at 3 dpf, larvae were tested at 6 dpf for escape behavior and prepulse inhibition. B. Increased SLC responses, decreased LLC responses and reduction in PPI after NTR-mediated ablation of *y252* neurons. Mann-Whitney U (MWU), * p < 0.001. C. Escape responses and PPI in control and photoablated dL5-expressing *y252* neurons. MWU, significant p-values indicated.

### Ablation of selected dL5-expressing neurons using spatially patterned illumination

Distinct neurons in the *y252-Gal4* pattern contribute to escape behavior and PPI. LLC responses require a cluster of *y252* neurons in the rhombomere 1 (R1) pontine tegmentum^17^ whereas PPI is mediated by glutamatergic neurons in rhombomere 4 (R4) of the medulla oblongata^16^. The location of neurons that regulate SLC thresholds are unknown. To test whether subsets of *y252* neurons could be selectively ablated, we used a digital micromirror device (DMD) to project spatially patterned far-red illumination onto defined brain regions (Fig. 3A). This approach resulted in selective ablation of either R1 or R4 *y252-Gal4, UAS:dL5-mCer* neurons (Fig. 3B-D). Consistent with our previous results using laser ablation, selective elimination of R1 *y252*-expressing neurons with light exposure left SLC and PPI responses intact while significantly reducing LLC responses (Fig. 3E). Conversely, SLC and LLC responses were not affected by selective R4 *y252* ablation, whereas PPI was substantially reduced (Fig. 3F). This double dissociation confirms that spatially restricted illumination can be used to selectively ablate dL5-expressing neurons of interest. Unlike NTR, that lesions the entire neuronal population defined by a transgenic pattern, spatially selective dL5-ablation addresses this limitation by enabling precise removal of neurons within a broad pattern, providing greater control over circuit manipulation for functional studies.

**Figure 3.**
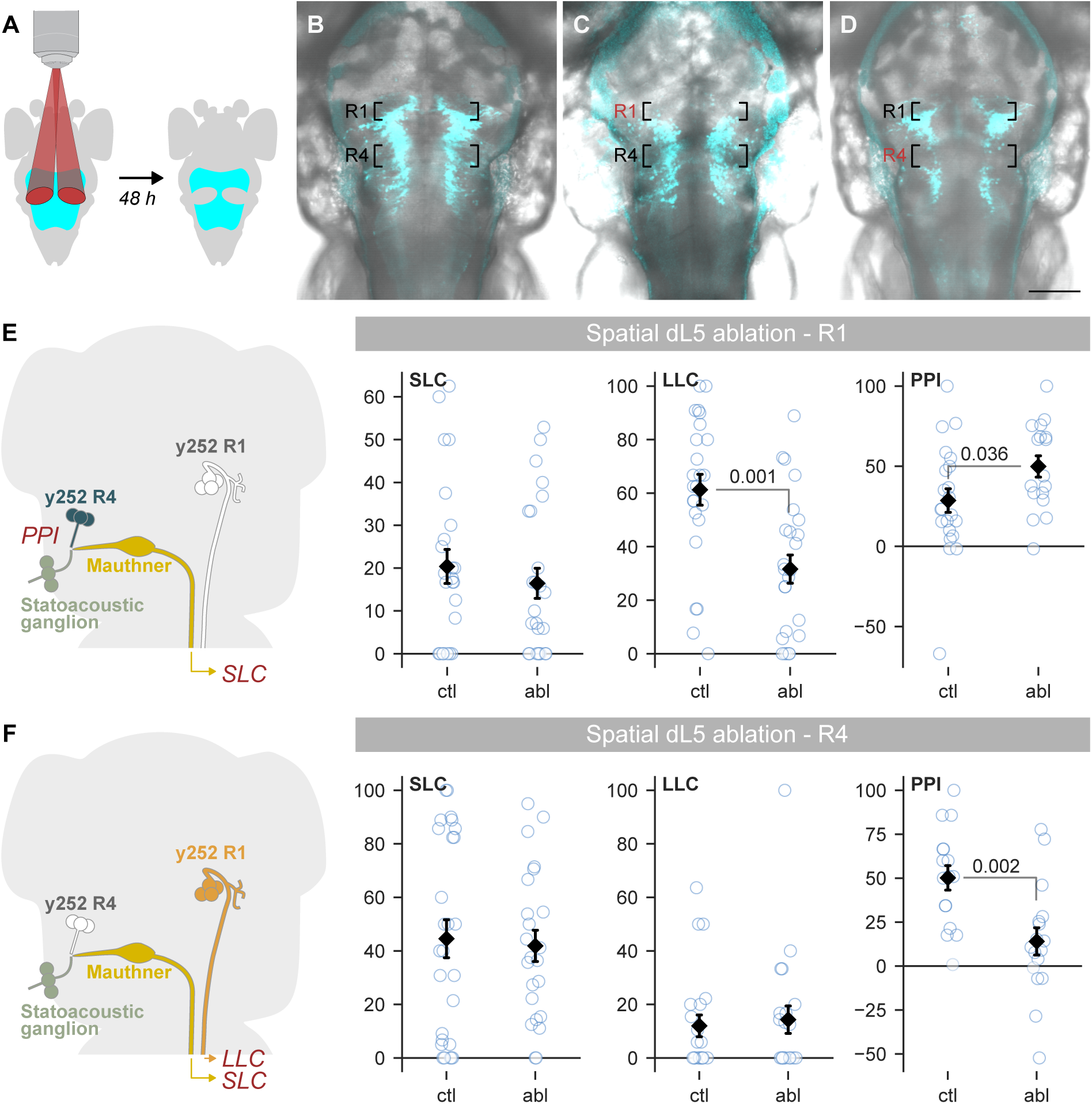
**Ablation of dL5 expressing neurons using spatially patterned illumination** A. Patterned illumination from a digital micromirror device used to selectively illuminate part of the *y252-Gal4* expression pattern. Larvae were tested for behavior 48 h after MG-2I treatment and photoablation. B-D. Confocal 40 µm substack maximum projections showing dL5-mCer expression and DIC background for 5 dpf untreated *y252-Gal4, UAS:dL5-mCer* larva (B), and larva 48 h after selective illumination of rhombomere 1 (C) or rhombomere 4 (D) neurons. Scale bar 100 µm. E-F. Escape behavior and prepulse inhibition in *y252-Gal4, UAS:dL5-mCer* larvae after selective photoablation of R1 neurons (E) or R4 neurons (F). MWU, significant p-values indicated. Increase in PPI in (E) was not present in replication experiment (ANOVA, main effect treatment F_1,54_ = 2.9, p = 0.094 for combined dataset).

### Presynaptically localized dL5 enables precise ablation of genetically defined synapses

Next, we constructed a synaptically localized version of dL5 through fusion to the vesicle protein synaptophysin^20^, and switched the fluorescent tag to mNeonGreen to improve our ability to resolve small patches of expression^21^(Fig. 4A). Like the dL5-mCer cassette, the synaptophysin-dL5- mNeonGreen (syp-dL5) cassette labeled cell bodies and neurites, but also showed punctate expression, consistent with synaptic localization (Fig. 4B,C). Immunostaining with the sv2 antibody, which labels *synaptic vesicle protein 2*, confirmed that puncta were presynaptic elements (Fig. 4B)^22^. Treatment of *y252-Gal4, UAS:syp-dL5* larvae with MG-2I and exposure to widefield illumination lead to a near complete loss of mNeonGreen fluorescence (Fig. 4C,D). When tested 72 h after light exposure, larvae reproduced the behavioral phenotypes generated by NTR ablation, including an increase in SLC responses, reduction in LLC responses and reduction in prepulse inhibition (Fig 4E). This shows that the syp-dL5 fusion protein retains phototoxicity.

**Figure 4.**
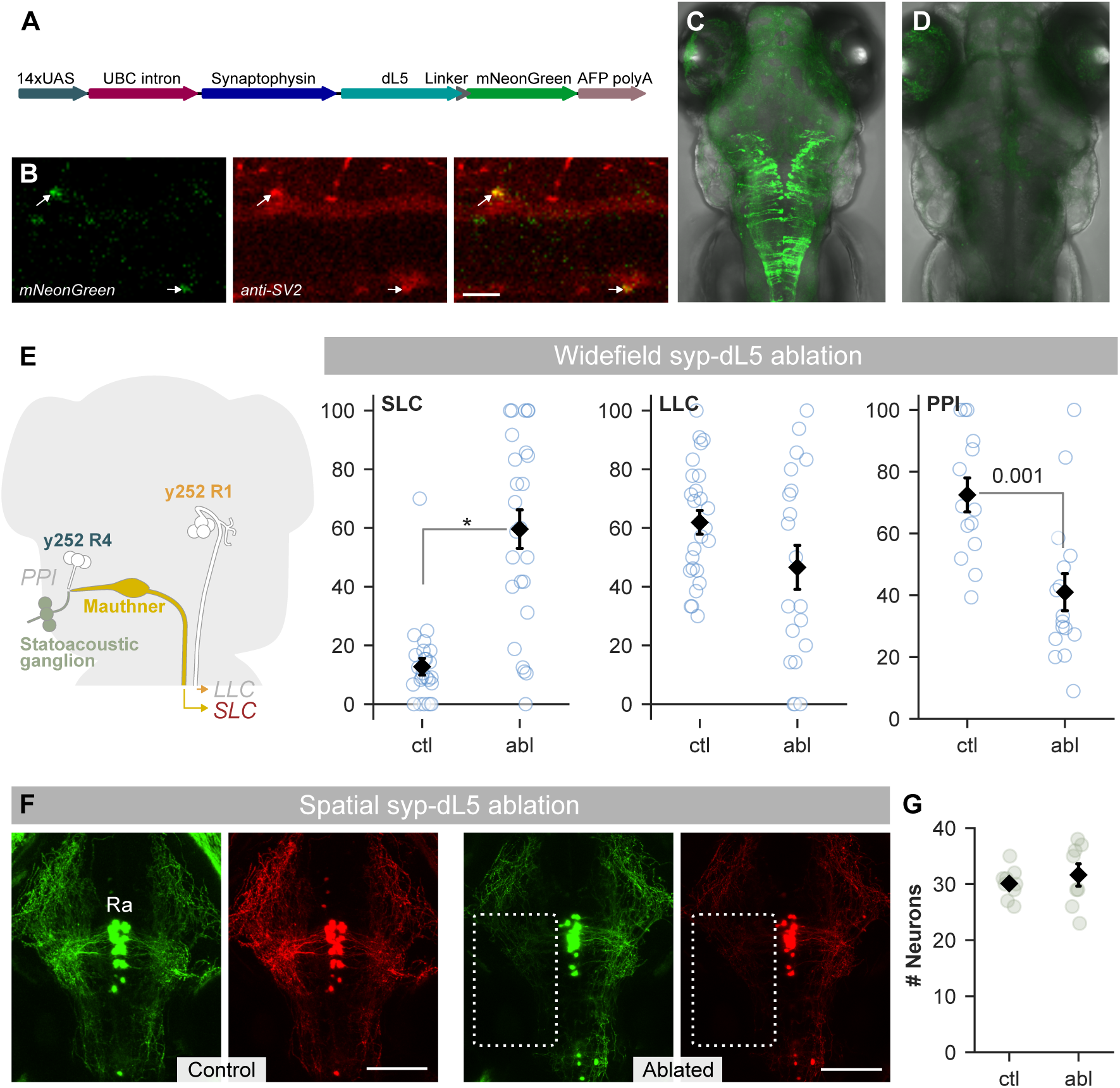
**Widefield ablation using synaptically localized dL5** A. Schematic of construct used to make *UAS:syp-dL5-mNeonGreen* transgenic line. B. Horizontal confocal section through the spinal cord of 3 dpf *y417-Gal4, UAS:syp-dL5-mNeonGreen* embryo (green), stained with anti-sv2 (red). Scale bar 20 µm. C-D. Maximum projection from confocal z-stack of GFP and DIC channels in 4 dpf *y252-Gal4, UAS:syp-dL5-mNeonGreen* larvae: untreated control larva (C) and an MG-2I treated larva exposed for 20 min to widefield illumination at 3 dpf (D). E. Escape responsiveness and prepulse inhibition in control and widefield light exposed *y252-Gal4, UAS:syp-dL5-mNeonGreen* larvae. MWU significant p-value indicated and * p < 0.001. F. Maximum projection of 80 µm confocal stacks from 4 dpf *tph2:Gal4, UAS:syp-dL5, UAS:Kaede^Red^* untreated larvae and in larvae exposed to NIR illumination in the indicated region of the left hindbrain at 3 dpf. Scale bar 100 µm. Ra, raphe neurons. G. Number of neurons in dorsal raphe 24 h after photoablation of neuropil in the region indicated in (F) in untreated (ctl) and photoablated larvae (abl). Welch t-test, p = 0.50

Because neuronal somas were also labeled by syp-dL5, it was possible that loss of the dL5 signal in the neuropil was secondary to photoablation of cell bodies. Conversely, we could not exclude the possibility that synaptic ablation leads to subsequent degeneration of cell bodies; this would preclude lesioning specific synapses to test their functional relevance. Finally, despite the lack of recovery over a 48 h period, we could not be certain that synapses were eliminated. Presynaptically localized miniSOG was reported to inactivate synapses by locally destroying adjacent vesicle release proteins, resulting in functional recovery after newly synthesized proteins were trafficked back to the synapse^9^. To assess whether syp-dL5 enabled selective lesioning of synapses we targeted a neuropil region innervated by raphe neurons in triple transgenic *tph2:Gal4, UAS:syp-dL5, UAS:Kaede* larvae. Because Kaede is cytoplasmic rather than vesicle-associated, we reasoned that if synapses were inactivated rather than ablated, most Kaede expression would persist even if dL5-mNeonGreen expression was lost. In addition, because raphe neurons are easily visualized, we could also determine whether synaptic ablation led to cell body degeneration. We first photoconverted Kaede from green to red with widefield UV light exposure, then mounted larvae for microscopy and used the DMD to train NIR light on a dense neuropil region in the left hindbrain innervated by raphe neurons (Fig. 4F). When we imaged larvae the next day, syp-dL5-mNeonGreen and Kaede^Red^ fluorescence were both almost completely eliminated in the selected region, suggesting that we had not merely photoablated local proteins but destroyed cellular components in the targeted region. Moreover, raphe neurons were intact, demonstrating that loss of synaptic label can be achieved without damage to cell bodies (Fig. 4G).

We previously showed that R4 *y252* neurons project to the lateral dendrite of the Mauthner cell, which receives input from statoacoustic ganglion (SAG) neuron afferents^16^. We found that during PPI, glutamate release from SAG neurons onto the Mauthner cell is depressed, consistent with presynaptic inhibition. Thus, ablating *y252* synapses in this region should disrupt PPI (Fig. 5A). To first test whether we could use patterned illumination to ablate synapses from *y252* neurons, we backfilled the Mauthner cells by injecting Alexa Fluor 658-dextran into the spinal cord in *y252-Gal4, UAS:syp-dL5* larvae. As expected, mNeonGreen puncta decorated the Mauthner cell lateral dendrite^15^. We then illuminated the lateral dendrite of only the left Mauthner cell. At 24 h, we imaged the Mauthner cells and saw a loss of mNeonGreen puncta apposed to the left Mauthner lateral dendrite (Fig. 5B). Thus, syp-dL5 expression can be used to ablate specific output synapses from genetically labeled neurons.

**Figure 5.**
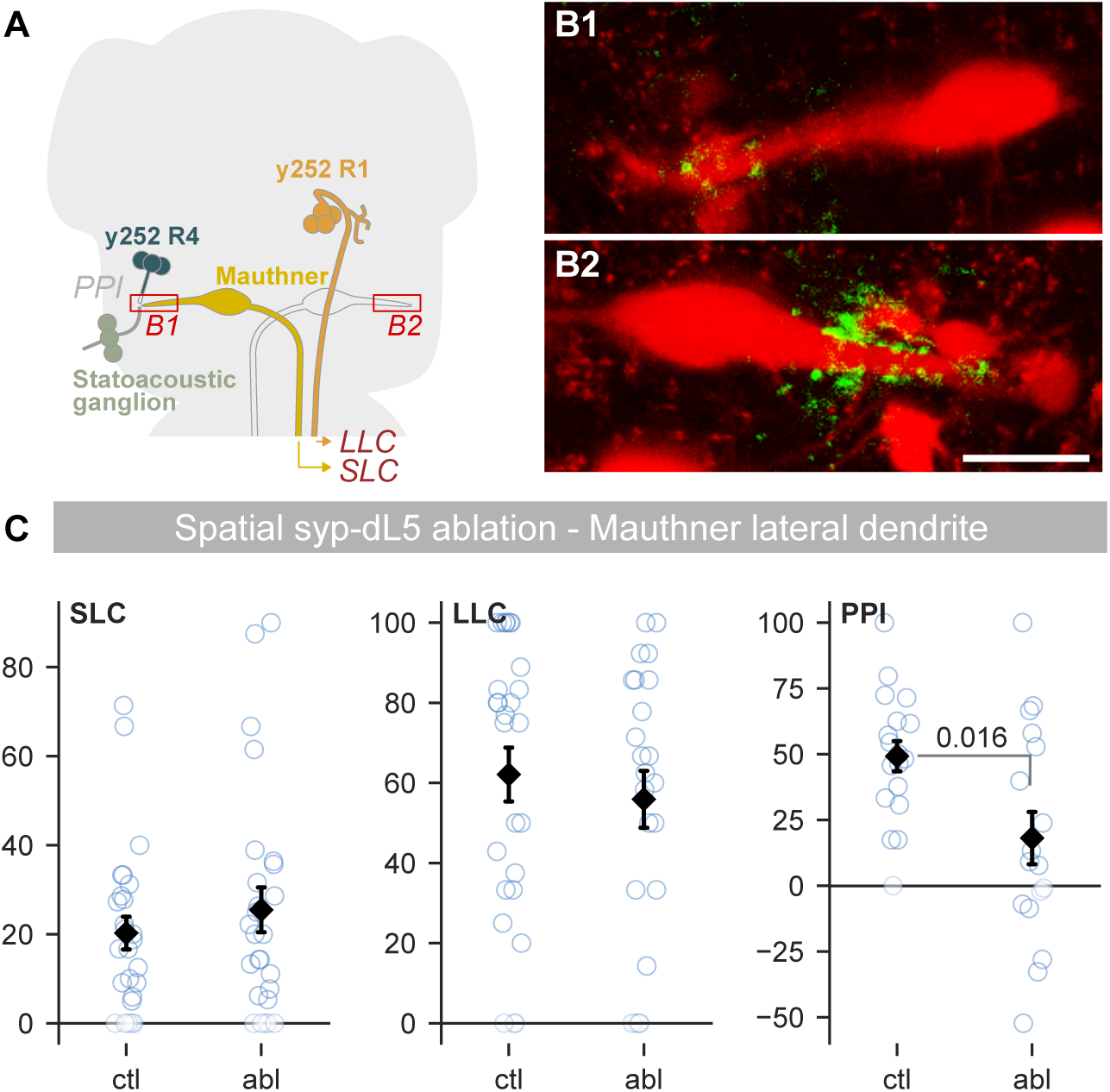
**Ablation of *y252* synapses at the Mauthner cell lateral dendrite impairs prepulse inhibition** A. *y252-Gal4* neurons project to the lateral dendrite of the Mauthner cell. Area of Mauthner cell dendrite illuminated in B1 indicated. Corresponding region in B2 was not light exposed. B. Rhodamine-dextran filled Mauthner cells (red) from an individual *y252-Gal4, UAS:syp-dL5* (green) larva, 24 h after illuminating the lateral dendrite of the left (B1) but not right (B2) Mauthner cell. Scale bar 20 µm. C. Startle behavior after bilateral ablation of *y252-Gal4, UAS:syp-dL5* synapses at the Mauthner cell lateral dendrites. MWU, p-value indicated.

To assess whether synaptic photoablation effectively disrupted signaling, we crossed *y252-Gal4, UAS:syp-dL5* to *J1229*, a transgenic line that expresses GFP in the Mauthner cell, allowing us to visualize the lateral dendrite without injecting dye into the spinal cord^23^. Subsequently, we illuminated synapses on both left and right Mauthner cell lateral dendrites. In ablated larvae, SLC and LLC responses were unaffected, but PPI was decreased, consistent with a loss of synaptic signaling by *y252* neurons to this area (Fig. 5C). This supports the hypothesis that direct projections from *y252* neurons to this region mediate PPI, and demonstrates that synaptically localized dL5 expression can be used to disrupt signaling to specific postsynaptic partners.

## Discussion

A full mechanistic understanding of neuronal circuit function requires decoding critical transmission paths within the central nervous system. This is greatly complicated by the large number of input and output partners associated with any given neuron. Inactivation of selected output synapses from a genetically labeled neuron can help to resolve experimentally how specific neuronal connections contribute to circuit function. To achieve this, we employed dL5, a synthetic photosensitizer protein that, when localized to a variety of subcellular compartments, complexed with MG-2I and exposed to NIR, has been previously shown to efficiently ablate cells^12^. We first confirmed that cytoplasmic expression of dL5 in combination with patterned illumination efficiently killed neurons in a region of interest. Then, by fusing dL5 to the synaptic vesicle protein synaptophysin, we were able to lesion specific output synapses from labeled neurons. To validate this approach, we took advantage of the circuit model for prepulse inhibition of the zebrafish escape response, where we have a relatively advanced understanding of essential neurons, their connections and associated behaviors.

Synaptic ablation serves as a complement to optogenetic systems where light-sensitive proteins in axons or terminals can be photostimulated to trigger or suppress neurotransmitter release, providing a powerful method to assess the functional relevance of specific neuronal connections^4^. However, synaptically localized optogenetic inhibitors may in some cases be difficult to validate or even show counterintuitive consequences due to rebound responses or changes in vesicle release dynamics^4,8^, and direct photostimulation of presynaptically localized channelrhodopsin can drive neurotransmission but may also lead to synaptic depression^24^. Moreover, while implanted optic fibers can be used to selectively photostimulate terminals in mammals, they are infeasible for small experimental models like zebrafish. Three other systems allow for persistent synaptic inactivation. Illumination of synapses expressing optoSynC or opto-vTRAP components reversibly clusters vesicles, inhibiting release for around 15 or 30 min respectively^25,26^. These are elegant systems but the time window is too short to be useful in fish. A method that uses photoactivatable botulinum neurotoxin to cleave the neurotransmitter release protein VAMP2 provides rapid and persistent interference with signaling, but has significant background activity^27^. These systems are limited by their incomplete suppression of neurotransmission (by around 50%) and their reliance on blue light which has limited tissue penetration and restricts their utility in behavioral assays that require visual stimulation or at least illuminated conditions. We therefore developed a near infra-red (NIR) photosensitizer-based system to selectively ablate synapses from genetically labeled neurons enabling behavioral experiments in freely moving animals with a light source that does not disturb their visual function.

We based our system on earlier work in which presynaptic localization of miniSOG, a small flavin mononucleotide binding protein, efficiently inhibited synaptic transmission^9^. MiniSOG efficiently produces singlet oxygen, leading to the destruction of nearby synaptic release machinery. However, synapses inactivated using synaptic-miniSOG recovered within 24 h likely due to the trafficking of newly synthesized proteins, which constrains the window available for subsequent testing. In addition, miniSOG is activated by blue light, and flavin mononucleotide is naturally present in cells, raising the risk of ablation during under standard experimental conditions. In contrast, the dL5 photosensitizer produces singlet oxygen only during illumination with near NIR light in the presence of MG-2I ^12^. This enables us to raise larvae under normal conditions until they are treated for 24 h with MG-2I, during which time they remain in dim light. After exposure to NIR, MG-2I is washed out and larvae can be raised and tested under normal conditions. Ablation occurs over an approximately 24 h period after light exposure and persists for at least two days. This provides an ample window for larvae to recover from the procedure and be used for experiments.

Photosensitizers may be a valuable alternative to other methods for targeted cell ablation. We demonstrated that transgenic expression of cytoplasmic dL5 can be used to lesion cells in a region of interest using spatially patterned illumination. This is similar to a cellular ablation system based on mitochondrially-targeted expression of a HaloTag protein in combination with a rhodamine-based photosensitizer dye, which efficiently kills zebrafish neurons over a similar time span^28^. Neurons are commonly lesioned using a two-photon laser, a technique that provides very high spatial resolution but that is difficult to calibrate to avoid off-target damage to neighboring cells^29^. Intersectional genetic strategies can provide exquisitely selective ablation with little or no collateral damage but are limited by the availability of transgenic lines that label cells of interest^16,30,31^. On the other hand, photoablation using dL5 should also be carefully validated. Although we found that photoablated neurons did not cause death of adjacent cells, singlet oxygen generation is likely proportional to transgene expression levels, as well as light intensity and duration, and in some circumstances may be sufficient to cause bystander damage. Similarly, we could not determine whether diffusion of singlet oxygen from syp- dL5 labeled synapses also destroys neighboring termini. However, one observation suggests that synaptic loss is relatively selective. Auditory afferents from the statoacoustic ganglion contact the Mauthner cell lateral dendrite, in the same area as *y252* synapses. The finding that short latency C-starts were not affected after ablation of *y252* synapses adjacent to the Mauthner lateral dendrite therefore implies that there was not a substantial loss of auditory termini despite their close proximity.

An important characteristic of syp-dL5 based synaptic ablation in comparison to other optically controlled methods for synapse inactivation is that it is irreversible. Although we expect that eventually ablated synapses will be replaced, we have not seen restoration of function over at least a 3 day period. This persistent loss of synaptic connectivity is useful for behavioral studies, but limits application for experiments requiring within-subject controls. In addition, unlike other methods where illumination intensity can be used to tune levels of inhibition, synaptic ablation is intrinsically an all-or-nothing phenomenon. Nevertheless, synaptic ablation using syp-dL5 is well suited to the zebrafish system, where a large number of transgenic lines direct highly specific effector expression in neurons of interest, optical transparency provides experimental access to neurons throughout the brain, and MG-2I can simply be applied to bath water^14,32–35^. Synapse-specific inactivation may be adapted to other small model organisms where fiber implantation is infeasible, although fusion of dL5 to a synaptic vesicle protein other than synaptophysin may be needed^9^.

Our results show that presynaptically localized expression of the dL5 photosensitizer protein enables selective ablation of targeted synapses, leading to highly specific deficits in escape behavior. This technique should be easily adapted to almost any other circuit with genetically tractable neurons, and help to interrogate the function of unique synaptic connections.

## Methods

### Zebrafish husbandry

All experiments were approved by the National Institute of Child Health and Human Development Animal Care and Use Committee. Experiments were performed on larvae up to 7 dpf, before sex differentiation. Larvae were raised at 28°C on a 14:10 light dark cycle in E3 medium supplemented with 1.5 mM HEPES pH 7.3. Wild-type and transgenic larvae were maintained on a Tüpfel Longfin background acquired from the Zebrafish International Resource Center (ZIRC). Transgenic lines used were *y252Et* (*y252-Gal4*)^35^, *y417Et* (*y417-*Gal4)^36^b, j1229aGt (*j1229*)^23^*, Tg(tph2:Gal4)y228 (tph2:Gal4)*^37^, *Tg(UAS:Kaede)s1999t*(*UAS:Kaede*)^38^, *TgBAC(slc17a6b[vglut2a]:loxP-DsRed-loxP-GFP)nns14* (*vglut2a:DsRed*)^39^.

### Plasmid construction

To create plasmid UAS:dL5-mCer3, we subcloned the MBIC5-mCer3 cassette from pCS2+ MBIC5-mCer3 (gift from Michael Tsang, Addgene plasmid # 74116 ; http://n2t.net/addgene:74116 ; RRID:Addgene_74116)^12^) into pT1UciMP (Addgene plasmid # 62215 ; http://n2t.net/addgene:62215 ; RRID:Addgene_62215)^40^. To make a synaptically localized version of dL5, we first replaced mCer3 in UAS:dL5-mCer3 with zebrafish codon-optimized mNeonGreen^21,40^, then created a fusion with zebrafish *synaptophysin b*^20^, creating UAS:syp-dL5-mNeonGreen. Stable lines were generated by injecting plasmids with codon optimized tol1 transposase mRNA^40^.

### Ablations

Nitroreductase ablations used *Tg(UAS:epNTR-TagRFPT-utr.zb3)y362Tg* (*UAS:epNTR-RFP*), a variant of nitroreductase engineered for increased ablation efficiency^18^. Embryos were treated with 10 mM metronidazole (Sigma M1547) from 3 to 4 dpf under dim light conditions. Controls were siblings from the same clutch that lacked RFP expression, also treated with metronidazole. For dL5 ablations, the stock solution of MG-2I (gift from Marcel Bruchez, Carnegie Mellon University or custom synthesized by www.medchem101.com) was dissolved at 1 mM in ethanol and stored in aliquots at - 20°C. Stock was added to E3 medium to make a working solution of 750 nM, supplemented with 300 μM N-Phenylthiourea (PTU) to inhibit melanophore formation. The working solution was applied to 2 dpf larvae, which were then placed in a dark 28°C incubator. At 3 dpf, working solution was replaced by E3/PTU medium. For widefield ablation, 6-8 larvae were placed in a single chamber of a 48 well plate containing 1 mL E3. A 656 nm LED (UHP-T-650-EP, Prizmatix) coupled to a 3 mm light guide was used to illuminate the chamber for 20 min (or less for experiments testing dose-response), providing 550 mW/cm^2^ of power. After treatment, larvae were returned to a regular light-cycle incubator in fresh E3/PTU medium. Larvae were maintained in PTU for behavioral experiments so that we could subequently verify ablation using confocal imaging.

### Spatial ablation

After anesthesia with 0.2 mg/mL tricaine methanesulfonate (MS-222), larvae were transferred to 100 µL E3 medium mixed with 200 µL 2.5% low-melt agarose. Larvae were mounted in a 50 mm glass-bottom dish, covered in E3 medium and transferred to a microscope with a digital micromirror device (Mightex Polygon 400). The same LED used for widefield photoablation was coupled to the Polygon 400 using a 3 mm light guide and controlled via a National Instruments DAQ board. Light was focused on the sample using a 40×/0.8NA water immersion objective, and ROIs defined using Elements software (Nikon). The power delivered via the DMD was 511 mW/cm ^2^ and illumination lasted 20 min.

### Confocal imaging

Embryos were maintained in E3 medium with 300 μM PTU starting at 24 hours post-fertilization (hpf). For imaging at 3 or 4 dpf, larvae were anesthetized in MS-222 for 5 min and subsequently mounted in 2.5% low-melting-point agarose. For low-resolution imaging, larvae were positioned dorso-ventrally in 50 mm glass-bottom dishes (#1.5), and confocal image stacks were acquired using a laser-scanning confocal microscope (Nikon AXR) equipped with an automated stage and a 20×/1.0NA objective. For high-resolution imaging, larvae were mounted in 3D-printed plastic inserts, which were placed on Lab-Tak II chambered cover glasses (#1.5). Confocal stacks were obtained using either a laser-scanning confocal microscope (Leica TCS SP5 II) with a 40×/0.95NA water immersion objective, or a point scan Nikon microscope with a 40×/1.15 NA objective.

### Histological labeling

To label the Mauthner cell with fluorescent dye we pressure injected a solution of Alexa Fluor 568-conjugated dextran 10k, into the ventral spinal cord of 3 dpf *y252-Gal4, UAS:syp- dL5* larvae^41^. After 24 h, we mounted larvae for ablation using the DMD. To photoconvert Kaede from green to red, we exposed larvae to 405 nm ultraviolet light (Formlabs Form Cure) for 5 min. For whole mount immunohistochemical staining, 3 dpf zebrafish larvae were fixed in 4% paraformaldehyde (PFA) in phosphate buffer saline (PBS) at room temperature for 2 h. Samples were washed three times with PBS, then treated on ice with 0.25% Trypsin for permeabilization for 5 min, then blocked with 1% Blocking Reagent (Roche, Indianapolis, USA) that was prepared in PBS for 3 hours on a rocker at room temperature. Primary antibody incubation with mouse anti-SV-2 antibody (1:200, Developmental Studies Hybridoma Bank, DSHB, Iowa City, IA) in PBS-T (PBS, 1% Triton) for 2 days at 4°C. For the secondary antibody, samples were treated with goat anti-mouse Alexa Fluor 633 (1:500, Invitrogen Company, A-21052) in PBS-T overnight at 4°C.

### Microplate measurements

For quantification of Cerulean fluorescence intensity in *y252-Gal4, UAS:dL5-mCer* after photoablation, larvae were placed into a black walled, round bottom 96-well plate in medium with PTU and tricaine (one larva per well). For each timepoint, we recorded 10 readings per well over a 10 min period using a Gen 5 microplate reader (Biotek Instruments). Larvae were then returned to Petri dishes and incubated at 28°C until the next timepoint. Microplate reader filters were set at 430 nm for excitation and 491 nm for emission to detect mCer fluorescence. Data in figure 1C shows readings after subtraction of background values recorded from non-fluorescent larvae measured at the same timepoints.

### Behavioral analysis

To test auditory responses and prepulse inhibition, we used a National Instruments NI-DAQ card (PCI-6221) to generate 2 ms duration waveforms, delivered to larvae using a minishaker (4810, Bruel and Kjaer) as previously described^42^. We used a series of 80 stimuli that comprised a pseudorandom sequence of four types of stimuli separated by 15 s. Stimuli were a weak vibrational stimulus, strong stimulus, weak stimulus with a 50 ms interval before the strong stimulus (PPI-50), and weak stimulus with a 500 ms interval before the strong stimulus (PPI-500). The weak stimulus was 10% of the intensity of the strong stimulus. Prepulse inhibition was calculated as the percentage reduction in short latency C-start responsiveness on PPI trials compared to responsiveness on trials with the strong stimulus alone. For experiments with *y252*, we report (1) short-latency C-start responsiveness to the weak stimulus, which increases after ablation, (2) long-latency C-start responsiveness, which decreases after ablation, and (3) prepulse inhibition from the PPI-500 measurement, which also decreases after ablation^15^. In some experiments, ablation of *y252* neurons with NTR caused an increase in PPI-50, but this effect was inconsistent between experiments, was not seen with dL5 and the underlying neural mechanism is unknown. Responses were recorded at 1000 frames per second using a high-speed camera then tracked and analyzed using Flote software^42^.

### Statistics

Data was analyzed using Python 3.8 and SciPy^43^. For behavioral experiments comparing startle responsiveness (SLC and LLC) and prepulse inhibition (PPI) between control and ablated larvae, values were generally not normally distributed. We therefore compared groups using a Mann-Whitney U (MWU) test. We used alpha 0.05 as the significance threshold. In plots of behavioral data, black diamonds and error bars represent the mean and standard error. We provide exact p-values in graphs except where p < 0.001, which are indicated using an asterisk.

## Acknowledgments

This work was supported by the Intramural Research Program of the *Eunice Kennedy Shriver* National Institute for Child Health and Human Development (NICHD) and utilized the high-performance computational capabilities of the Biowulf Linux cluster at the National Institutes of Health, Bethesda, MD. We thank Jennifer Panlilio for her valuable contributions. We thank Leanne Iannucci for help calibrating the DMD. EAB acknowledges support from the US Department of Veterans Affairs (BX003168), NIH (NS125280) and the Parkinson Foundation (PF-IMP-1259272). The contents of this article do not represent the views of the United States Government.

