## Supplementary figures and images for "Functional interrogation of neuronal connections by chemoptogenetic presynaptic ablation"

### Supplementary Figure 1

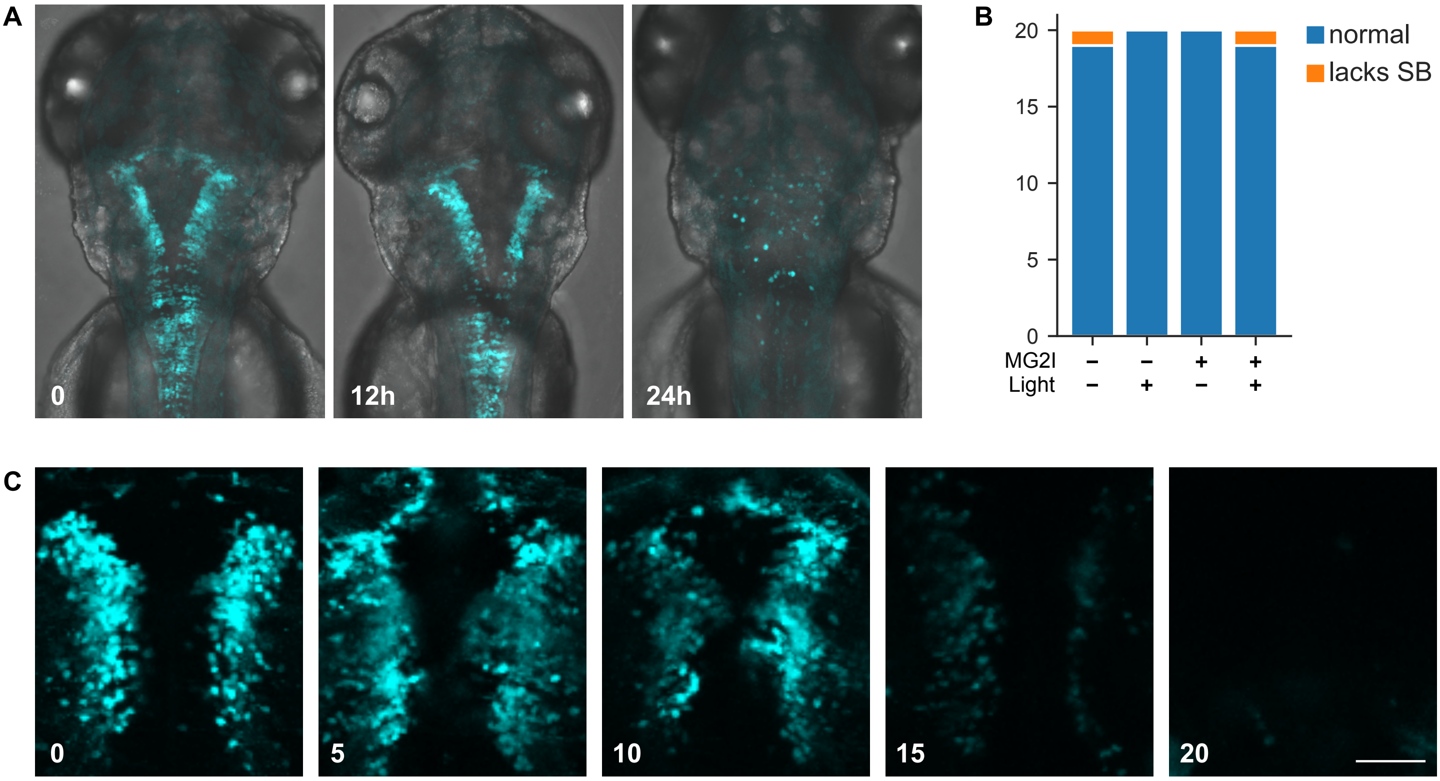


Supplementary Figure 1
